# 2½-minute 3D 7T ^31^P-MRSI of the human heart using concentric rings (CRT)

**DOI:** 10.1101/2021.12.10.472120

**Authors:** William Clarke, Lukas Hingerl, Bernhard Strasser, Wolfgang Bogner, Ladislav Valkovič, Christopher T Rodgers

**Affiliations:** Wellcome Centre for Integrative Neuroimaging, FMRIB, Nuffield Department of Clinical Neurosciences, University of Oxford, Oxford, UK; High-field MR Centre, Department of Biomedical Imaging and Image-guided Therapy, Medical University of Vienna, Vienna, Austria; Oxford Centre for Clinical Magnetic Resonance Research, Radcliffe Department of Medicine, University of Oxford, Oxford, UK; Department of Imaging Methods, Institute of Measurement Science, Slovak Academy of Sciences, Bratislava, Slovakia; Wolfson Brain Imaging Centre, Department of Clinical Neurosciences, University of Cambridge, Cambridge, United Kingdom

**Keywords:** Spectroscopy, ^31^P, phosphorus, MRSI, heart, CRT

## Abstract

A 3D density-weighted concentric ring trajectory (CRT) MRSI sequence is implemented for cardiac ^31^P-MRS at 7T.

The point-by-point k-space sampling of traditional phase-encoded CSI sequences severely restricts the minimum scan time at higher spatial resolutions. Our proposed CRT sequence implements a stack of concentric rings trajectory, with a variable number of rings and planes spaced to optimise the density of k-space weighting. This creates flexibility in acquisition time, allowing acquisitions substantially faster than traditional phase-encoded CSI sequences, while retaining high SNR.

We first characterise the signal-to-noise ratio and point spread function of the CRT sequence in phantoms. We then evaluate it at five different acquisition times and spatial resolutions in the hearts of five healthy participants at 7T. These different sequence durations are compared with existing published 3D acquisition-weighted CSI sequences with matched acquisition times and spatial resolutions. To minimise the effect of noise on the short acquisitions, low-rank denoising of the spatio-temporal data was also performed after acquisition.

The proposed sequence measures 3D localised PCr/ATP ratios of the human myocardium in 2.5 minutes, 2.6 times faster than the minimum scan time for the acquisition-weighted phase-encoded CSI. Alternatively, in the same scan time a 1.7-times smaller nominal voxel volume can be achieved. Low-rank denoising reduced the variance of measured PCr/ATP ratios by 11% across all protocols. The faster acquisitions permitted by 7T CRT ^31^P-MRSI could make cardiac stress protocols or creatine kinase rate measurements (which involve repeated scans) more tolerable for patients without sacrificing spatial resolution.

## Introduction

Phosphorus magnetic resonance spectroscopy (^31^P-MRS) allows measurement of the energy metabolism of the human heart *in vivo*, specifically the ratio of phosphocreatine to adenosine triphosphate (PCr/ATP), which is a biomarker of heart failure (1). To date, 3D-localized ^31^P-MRS measurements of the human heart have used chemical shift imaging (CSI) with Cartesian phase-encoded k-space sampling or single voxel 3D-image-selected in vivo spectroscopy (ISIS) pulse sequences (2,3). While CSI offers optimal SNR per unit time, the point-by-point sampling of k-space severely restricts the minimum scan time at higher spatial resolutions (4).

Long acquisition times can restrict our ability to acquire data in a timeframe that is tolerable for clinical purposes. However, they are particularly restrictive when multiple, repeated acquisitions are needed for either stress protocols (5), or to non-invasively measure chemical kinetics of the oxidative phosphorylation system (6,7). These protocols which are thought to provide more sensitive detection of underlying pathological processes (8,9) have typically been achieved by lowering spatial resolution, leading to significant partial volume effects (10,11).

Employing fast MRSI readout trajectories, it could be possible to leverage the approximately 2.8 times increase in SNR (3) to achieve close to the theoretical 7.8-times (2.8^2^) speed increase when moving from 3T to 7T, which is not feasible with point-by-point Cartesian sampling. Concentric ring trajectory (CRT)-MRSI is an attractive option because it has been shown to deliver high-resolution ^1^H-MRSI of the brain with close to optimal SNR-per-unit-time (12,13).

Here we propose a 3D density-weighted CRT-MRSI sequence to achieve fast ^31^P-MRSI of the human heart. This is achieved by modifying a previously implemented ^1^H-CRT-MRSI sequence (14) to include full 3D density weighting to achieve a compact three dimensional point spread function in the acquisition, thereby avoiding the loss of SNR associated with post-acquisition reweighting (15).

MRSI acquisitions have highly redundant data, and are therefore particularly well suited to post-processing with low-rank denoising (16,17) to improve metabolite quantification precision. Here low-rank denoising could mitigate the expected loss of SNR when reducing the sequence acquisition time. We therefore also compared the effects of an optimised low-rank denoising approach (17) on data acquired using CSI and CRT trajectories.

In this work we demonstrate the feasibility of a 3D density-weighted CRT sequence for rapid ^31^P-MRSI of the human heart at 7T. We compare this methodology to previously published sequences for reduced acquisition duration (with fixed resolution) or increased spatial resolution (with equivalent maximum scan time) (11). In addition, we assess the impact of modern optimised low-rank denoising (17) postprocessing on rapidly acquired CSI and CRT MRSI data. We aim ultimately to present a state-of-the-art approach to human cardiac metabolic imaging.

## Methods

### Sequence design

A density-weighted 3D-CRT sequence was created by modifying a previously published equidistant ring 3D-CRT sequence (14). The sequence diagram of the modified sequence is shown in Figure 1. The MUSICAL (18) coil-sensitivity scans were removed from the original sequence because they rely on the unsuppressed water signal which has no analogue in ^31^P-MRSI. The CRT readout gradients were modified with 3D density weighting (Figure 2). The density weighting function *w*(*k*) was implemented as described in Equation 1.^1^

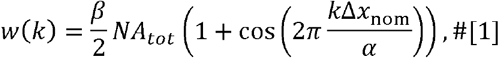

where Δ*x*nom is the nominal spatial resolution, and *NA*_*tot*_ is the total number of acquisitions which for this work is set to one. *α* and *β* are constants set according to the Raleigh criterion, as described elsewhere(15,19). The weighting was implemented by placing rings with irregularly spaced radii (in the k_xy_-plane) on irregularly spaced planes (in the k_z_-direction, Figure 2). In this work *α* was set to 1.71 in the k_xy_-plane and 1.61 in the k_z_-direction. *β*was set to 1.47 and 1.25 respectively. *α* and *β* values were chosen from literature values (15,19) and simulation of the PSF for the CRT sequence trajectory with the 1D, 2D, and 3D *α*and *β*values given in the literature. Density weighting in the k_xy_ plane (concentric rings) was implemented in the sequence using the process described in Reference (13). For the k_z_ direction the position of the planes was calculated similarly; plane positions were calculated by uniformly sampling along the cumulative distribution function of Equation 1. This was implemented by using a series expansion to numerically approximate the inverse cumulative distribution function. A detailed description of the implementation is provided in the supporting information.

**Figure 1.**
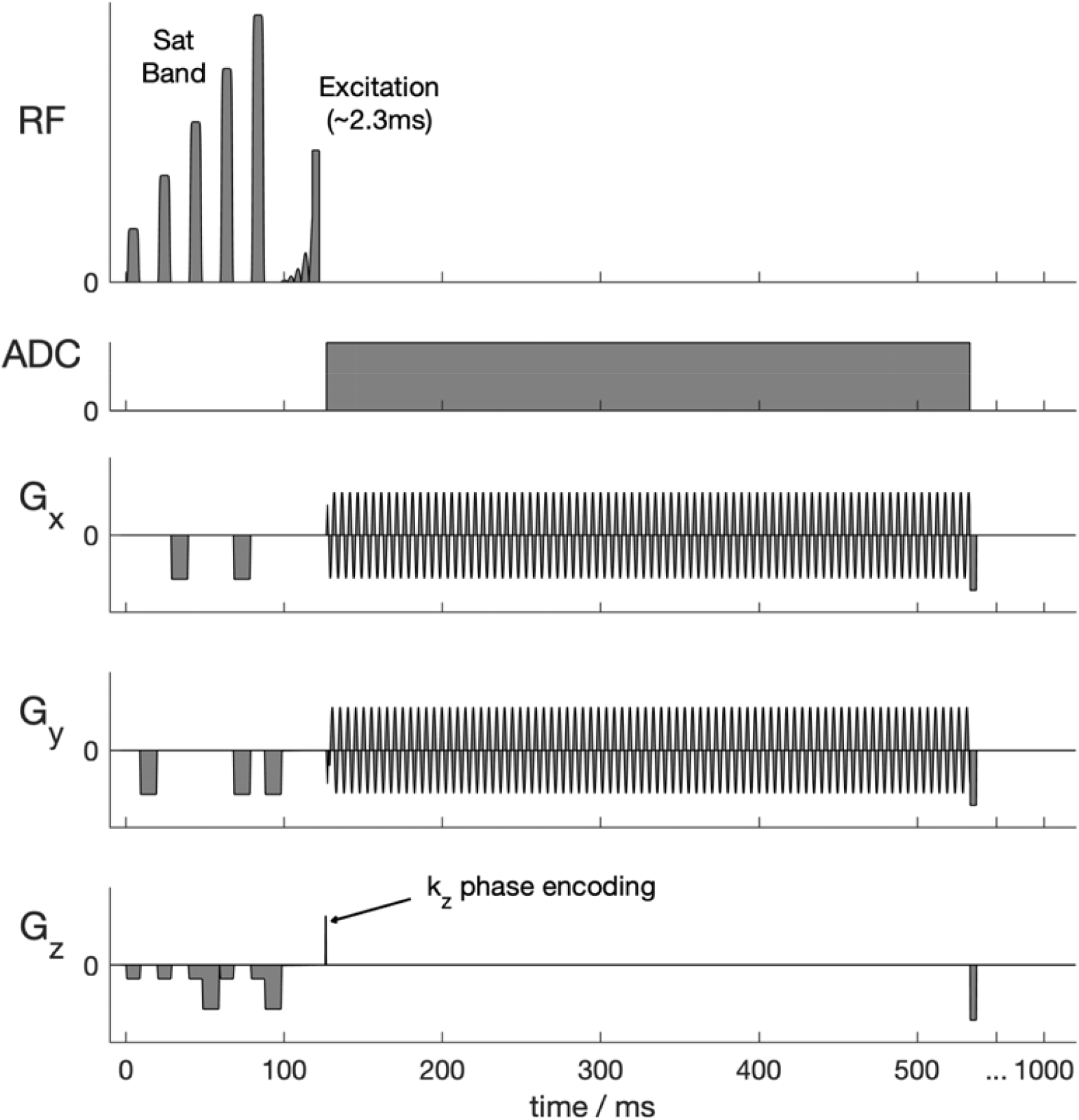
Pulse sequence diagram (single repetition time). Not to scale. The 3D-CRT sequence with density weighted k-space acquisition is preceded by a BISTRO saturation band module suitable for suppressing skeletal muscle signal at 7T using surface coils. An asymmetric shaped excitation pulse providing minimal amplitude and phase variation over a ∼2.5 kHz bandwidth was implemented (Figure 4 of Reference (3)).

**Figure 2.**
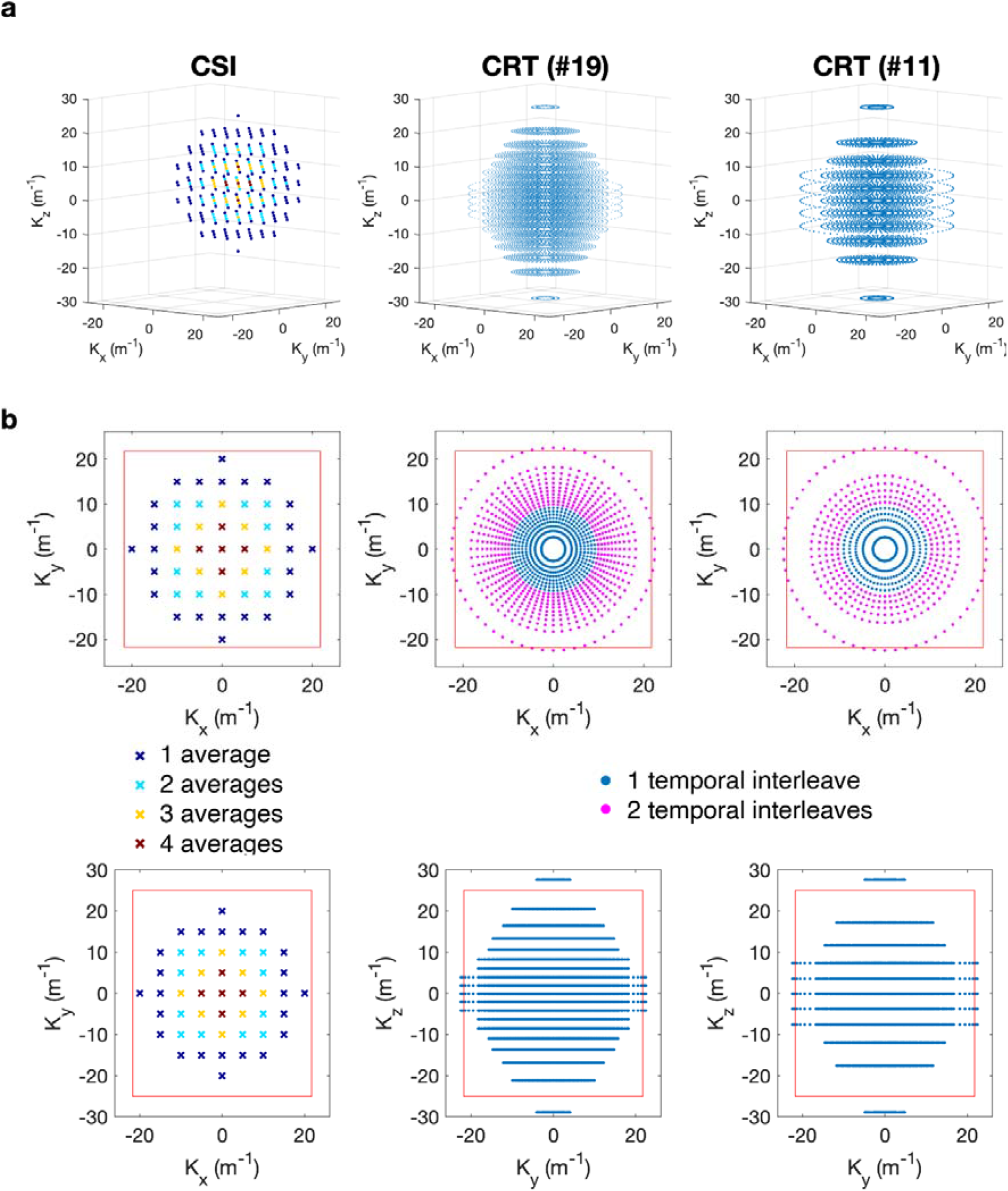
**a** 3D plots of the k-space trajectories of the Cartesian acquisition weighted CSI (left), CRT sequence with 19 rings/partitions (middle) and CRT sequence with 11 rings/partitions (right). At each k-space location marked a time domain signal is acquired (not shown). **b** Trajectories of the sequences shown in the xy-plane (top) and yz-plane (bottom). The red box marks the extent of the k-space (1/Δ_x/y_). The CSI acquisition weighting is illustrated via different colour-coding, with the central points sampled more frequently than the outer. The magenta points in the CRT sequence show rings acquired using two temporal interleaves to overcome spectral bandwidth limitations.

To complement the density weighting, the number of rings in each plane was tuned to elliptically sample k-space in the k_z_-direction. Up to two temporal interleaves were used to achieve a fixed dwell time (spectral bandwidth) in rings that would otherwise have exceeded the hardware gradient slew rate limits. Total sequence duration was adjusted by varying the number of rings and k_z_ partitions whilst keeping the maximum k-space coverage identical and thus the nominal resolution in the x, y, and z directions equal.

To aid comparison with the established CSI sequence, we have implemented the same RF excitation pulse and outer volume saturation scheme as previously described (3). The excitation used an asymmetric 2.4 ms, shaped, constant phase pulse (Figure 1) designed to uniformly excite metabolites between -3 and 8 ppm (i.e., 2,3-DPG, PDE, PCr and *γ*-ATP) when centred 270 Hz from PCr. A single “B_1_-insensitive train to obliterate signal” (BISTRO) saturation band to suppress the chest wall signal was added to the sequence (20), and applied each repetition time.

### Reconstruction

CSI data were reconstructed online using a modified version of the vendor’s reconstruction code (3,21). CRT data were reconstructed offline using the non-uniform FFT (NUFFT) toolbox with min-max Kaiser-Bessel kernel interpolation and twofold oversampling (22) in MATLAB (MathWorks, Natick, USA). Density compensation was not applied in addition to the trajectory density weighting. Individual coil data was combined using the WSVD algorithm (21).

### Simulation and phantom validation

The density-weighted CRT sequence was validated through simulations and phantom scans. The SNR and point-spread function (PSF) of the sequence was characterised relative to an acquisition weighted CSI sequence on a point-source phantom (3). The measured PSF was compared to the numerically simulated PSF.

CSI and CRT data with closely matched parameters were acquired on a phantom containing a 2×2×2 cm^3^ cube of 1 M K_2_ HPO_4_ in a large tank filled with saline. The acquisition grid was placed to centre a voxel over the point source. Seven different acquisitions were made with matched field of view (200×200×200 mm^3^), spectral bandwidth (8 kHz), T_R_ (1 s), and RF pulse voltages:

1. Acquisition-weighted CSI with 2×2×2 cm^3^ resolution, 10×10×10 matrix and N=4 at k=0 giving TA of 6:31 min;
2. As #1 but with N=1 at k=0 (TA: 4:31 min);
3. CRT with 19, 15, 13, and 11 rings/partitions (TA: 6:55, 4:12, 3:12, and 2:18 mins) reconstructed to a 10×10×10 Nyquist matrix;
4. CRT with 18 rings/partitions (6:27 mins) reconstructed to a Nyquist matrix size of 12×12×12.

The PSF was predicted by passing a uniform synthetic signal through the NUFFT adjoint operation, as formulated for the expected gradient trajectory. The predicted SNR was calculated accounting for the total acquisition time and voxel volume and normalised to the SNR measured for the acquisition weighted CSI acquisition to account for effects such as phantom T_1_, coil sensitivity that are equal in all scans. Phantom measured PSF and SNR were compared to predicted values.

### In vivo comparison of CSI and CRT

Five healthy subjects (4 male & 1 female, 74±11 kg, 30±3 years) were scanned in a supine position using a whole-body Siemens Magnetom 7T scanner (Erlangen, Germany) equipped with a combined 10 cm ^1^H / 15 cm ^31^P quadrature-pair transmit-receive surface coil positioned over the heart (23). Each subject was scanned using a range of CSI and CRT sequences with different acquisition times and sampling densities. The specific details of each sequence are given in Table 1. The protocols included a previously described CSI sequence (11) (Protocol 1 in Table 1). Hereafter, we refer to each protocol as 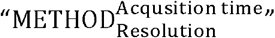, for example, 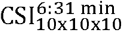 is Protocol 2 in Table 1. Other protocol parameters were closely matched to those used for the previously described CSI sequence. Position, orientation, field of view (FoV), RF pulse voltages, and repetition times of all protocols were matched. Readout bandwidth was dependent on acquisition method because the CRT spectral bandwidth is limited by hardware gradient slew rates. A maximum of two temporal interleaves were used to not prolong acquisition time. This resulted in a CRT bandwidth less than CSI, which has an excessive bandwidth (8000 Hz) for the excited bandwidth (approximately 2500 Hz).

**Table 1.** Top: Protocol parameters. Colour coding is matched to subsequent figures. Bottom: Results. Matched-filter PCr SNR, mean(±SD) uncorrected PCr/ATP, saturation corrected PCr/ATP, and number of voxels fitted with PCr/ATP CRLB <30% from the septal voxels (total 52, 10-11 voxels per subject). Mean and SD is calculated over all septal voxels of all subjects. The table also shows the matched-filter PCr SNR and saturation corrected PCr/ATP values from the same voxels after denoising. “SNR Ratio” is the ratio of “Denoised PCr SNR” to “PCr SNR”.

| Protocol # | 1 | 2 | 3 | 4 | 5 | 6 | 7 | 8 |
| --- | --- | --- | --- | --- | --- | --- | --- | --- |
| **Acquisition parameters** |  |  |  |  |  |  |  |  |
| **Trajectory** | CSI | CSI | CSI | CRT | CRT | CRT | CRT | CRT |
| **TA (m:ss)** | 6:37 | 6:31 | 4:21 | 6:32 | 4:33 | 3:19 | 2:31 | 6:55 |
| **NA (k=0)** | 4 | 4 | 1 |  |  |  |  |  |
| **# rings/partitions** |  |  |  | 20 | 16 | 14 | 12 | 19 |
| **Reconstruction matrix** | 8x16x8 | 10x10x10 | 10x10x10 | 10x10x10 | 10x10x10 | 10x10x10 | 10x10x10 | 12x12x12 |
| **Nominal voxel (ml)** | 11.3 | 11.5 | 11.5 | 11.5 | 11.5 | 11.5 | 11.5 | 6.7 |
| **FoV (mm<sup>3</sup>)** | 240*240*200 | 240*240*200 | 240*240*200 | 240*240*200 | 240*240*200 | 240*240*200 | 240*240*200 | 240*240*200 |
| **T<sub>R</sub> (s)** | 1 | 1 | 1 | 1 | 1 | 1 | 1 | 1 |
| **Bandwidth (Hz)** | 8000 | 8000 | 8000 | 2778 | 2778 | 2778 | 2778 | 2778 |
| **Time samples** | 2048 | 2048 | 2048 | 720 | 720 | 720 | 720 | 720 |
| **TA reduction compared to #1** | 1.00 | 1.02 | 1.52 | 1.01 | 1.45 | 1.99 | 2.63 | 0.96 |
| **Results** |  |  |  |  |  |  |  |  |
| **PCr SNR** | 14.5 (±10.9) | 15.2 (±8.8) | 11.9 (±7.2) | 14.9 (±10.6) | 10.9 (±7.2) | 9.9 (±7.3) | 9.0 (±6.0) | 9.2 (±6.8) |
| **PCr/ATP** | 1.07 (±0.63) | 1.21 (±0.52) | 1.37 (±0.59) | 1.22 (±0.56) | 1.19 (±0.58) | 1.14 (±0.87) | 1.03 (±0.71) | 1.31 (±0.53) |
| **Corrected PCr/ATP** | 1.85 (±0.89) | 1.90 (±0.93) | 1.82 (±0.97) | 1.71 (±0.96) | 1.80 (±1.02) | 1.59 (±0.93) | 1.52 (±0.97) | 2.12 (±0.79) |
| **Septal voxels (#/52)** |  |  |  |  |  |  |  |  |
| **CRLB <30%** | 49 | 52 | 48 | 49 | 47 | 43 | 43 | 45 |
| **CRLB <25%** | 47 | 50 | 47 | 48 | 44 | 40 | 38 | 39 |
| **Denoised PCr SNR** | 31.3 (±15.6) | 40.4 (±15.5) | 24.6 (±9.9) | 26.8 (±12.6) | 20.8 (±9.2) | 21.7 (±10.7) | 19.7 (±9.3) | 20.8 (±10.2) |
| **SNR Ratio** | 2.2 | 2.7 | 2.1 | 1.8 | 1.9 | 2.2 | 2.2 | 2.3 |
| **Denoised Corrected PCr/ATP** | 1.80 (±0.81) | 2.01 (±0.74) | 1.91 (±0.79) | 1.94 (±0.89) | 1.93 (±0.87) | 1.95 (±0.95) | 1.57 (±0.95) | 2.08 (±0.95) |

Additionally, ^1^H CINE FLASH images with pulse oximeter gating were acquired in each subject for anatomical spectral localisation and VOI identification. Images were acquired in 4-chamber and 2-chamber long axis and three short axis views (apical, mid, and basal). From these structural images anterior-, mid-, and posterior interventricular septal voxels from apical, mid, and basal short-axis views of the heart were manually picked based on our institution’s standard anatomical landmarks. In addition, pure ventricular voxels (blood voxels) were selected from the right and left ventricles from apical, mid, and basal short-axis views. Voxels were selected based on the 10×10×10 CSI grid with nearest neighbour interpolation used to select voxels from other resolutions. Selection was done blinded to the MRS data.

### Pre-processing and fitting

Data processing was carried out using the OXSA toolbox (24). Spectra were corrected for frequency offset and DC offset. Then peaks were fit with the “Advanced method” for accurate, robust and efficient spectral fitting (AMARES) (24,25). Prior knowledge specified 11 Lorentzian peaks (α,β,ATP multiplet components, PCr, PDE, and the two peaks of 2,3-diphosphoglyceric acid [DPG]), with fixed amplitude ratios and scalar couplings for each multiplet. Fitting was initialised with begin times measured from each sequence simulated in the vendor’s simulation environment, corresponding to of 0.35 ms and 0.85 ms for CSI and CRT, respectively. Blood contamination and partial saturation were corrected as previously described (3,24). Metabolite ratios and ratio uncertainties are reported for PCr/ATP.

To aid sequence comparison irrespective of inter-subject differences in PCr/ATP ratio, we also computed normalised PCr/ATP ratios according to:

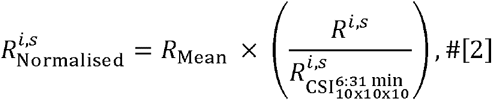

Where is the saturation and blood corrected PCr/ATP ratio of the *i*^th^ voxel of the *s*^th^ subject for a particular protocol. scaled all values to the mean saturation and blood corrected PCr/ATP value of the protocol.

### Spatio-temporal denoising

Local (sliding-window) low-rank spatio-temporal denoising was applied to all reconstructed frequency domain MRSI data (16,17). Patch size was chosen to be 3×3×3 with a stride of one in all directions. Automatic rank selection was applied patch-wise using the Marchenko-Pastur distribution method (17,26). This formed a denoised representation of each reconstructed MRSI dataset. The denoising code is open-source and available online (https://git.fmrib.ox.ac.uk/wclarke/low-rank-denoising-tools), and as an installable package “mrs_denoising_tools” via the package managers PyPi (Python Software Foundation, Wilmington, DE, USA) and Conda (Anaconda Inc, Austin, TX, USA). The denoised data were also fitted and corrected for blood contamination and partial saturation, following the same procedure described above.

### Sequence comparison

Results were compared to the reference 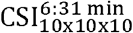 dataset using saturation and blood-corrected PCr/ATP ratios in the selected voxels using the Wilcoxon signed rank test (for paired measurements). The comparison was repeated for denoised results, while still comparing to the non-denoised reference 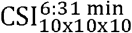 dataset.

## Results

### Simulation and phantom validation

Phantom experiments showed that the measured CRT PSF matched the predicted low ripple PSF both in the plane of the rings (Figure 3a&c) and through the plane in the third dimension (Figure 3b&d). The z-direction PSF of the CRT was narrower than the CSI potentially indicating a small deviation from the desired weighting function. Reducing the number of rings to 10 from 19 did not cause deterioration of the central lobe of the PSF (FWHM increased by 0.2%, Figure 3f). The high resolution, 18-ring 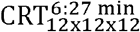 sequence had similar PSF with low ripple. The FWHM of the central lobe was 10% smaller than the 10×10×10 sequence, less than predicted by the resolution increase, but explained by the presence of signal arising outside the smaller voxel.

**Figure 3.**
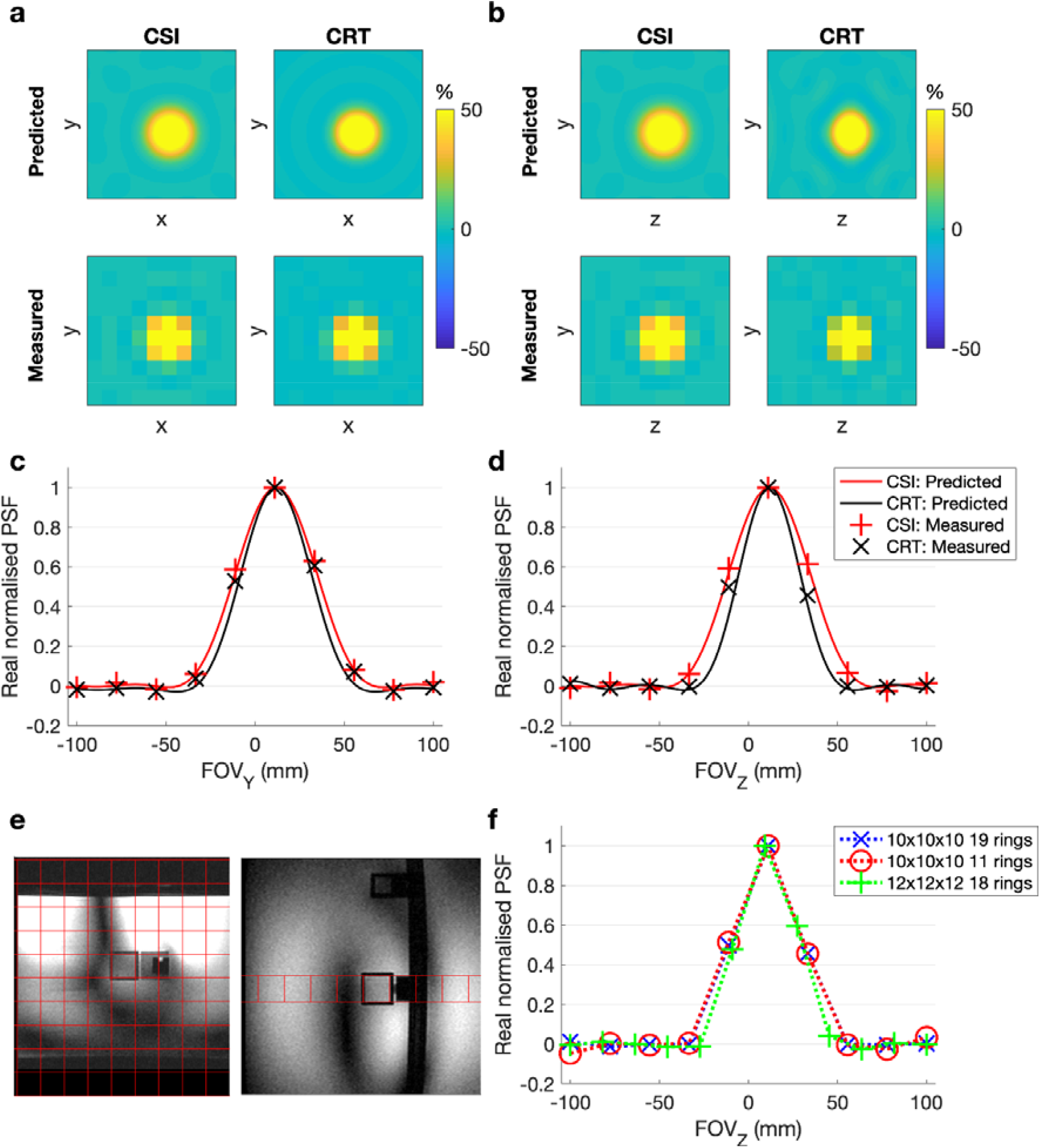
Predicted and measured point-spread-functions (PSF) of the CSI and 19 ring CRT sequence in the xy-plane (**a**) and yz-plane (**b**). The PSF profile in the y and z axis is plotted through the central point in **c** & **d. e** Point source phantom used to measure the PSF with the 10×10×10 spatial grid overlaid. ^31^P signal only arises from the central cube. **f** Effect of varying resolution and number of rings on the measured PSF in the z-direction.

SNR performance mostly showed the predicted relationship, dependent on acquisition time (Figure 4). SNR was matched between time-matched acquisition weighted CSI and density weighted CRT sequences. The post-acquisition reweighting of the single average (uniform weighted) CSI results in an SNR loss. Therefore the 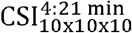 produced lower SNR than predicted, and lower SNR than the density-weighted 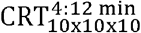 scan. The high-resolution 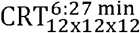 produced higher SNR than predicted, but this is likely to arise from the signal outside the measured voxel bleeding into the measured voxel because of the PSF.

**Figure 4.**
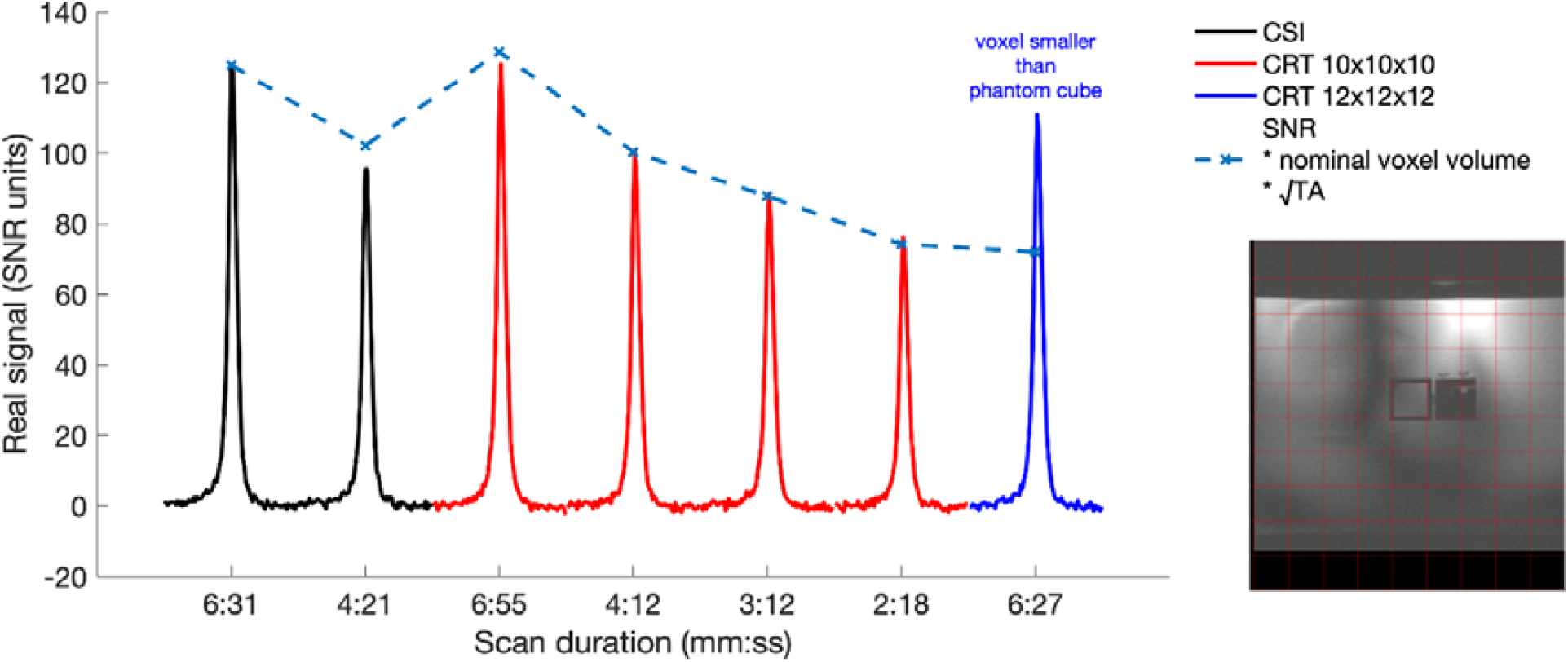
Measured signal (scaled to SNR units) from the point source phantom acquired with CSI and CRT sequences of varying acquisition times. The predicted SNR is calculated relative to the 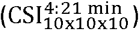 scan, adjusting for the nominal voxel volume and acquisition time of each protocol. The deviation for the short 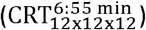 scan arises because CSI can only sample integer numbers of points at each k-space location, thus SNR is lost in post-acquisition reweighting. The larger deviation for the higher resolution CRT scan (6:27 min) has a voxel smaller than the ‘point-source’ resulting in signal bleed. The image shows the position of the 10×10 imaging matrix (red) over the phantom ‘point-source’. ^31^P signal only arises from the central cube.

### In vivo results

^31^P-MRSI was acquired successfully using 3D density-weighted CRT with acquisition times down to 2:31 in all five subjects. Example PCr/ATP and PCr SNR maps of the mid-short-axis slice from four sequences in one subject are shown in Figure 5a. Maps for all sequences are shown in supporting information Figure S1. The selection of voxels based on standard anatomical landmarks to include only cardiac and surrounding voxels resulted in 52 septal myocardial voxels selected across five subjects (10-11 per subject; three apical, three or four mid, and four basal), and 10 ventricular voxels (two per subject). Example manually selected inter-ventricular septal, and ventricular voxels for analysis are shown in Figure 5b. Spectra from mid-septal voxels of the same four sequences and subject are shown in Figure 5c.

**Figure 5.**
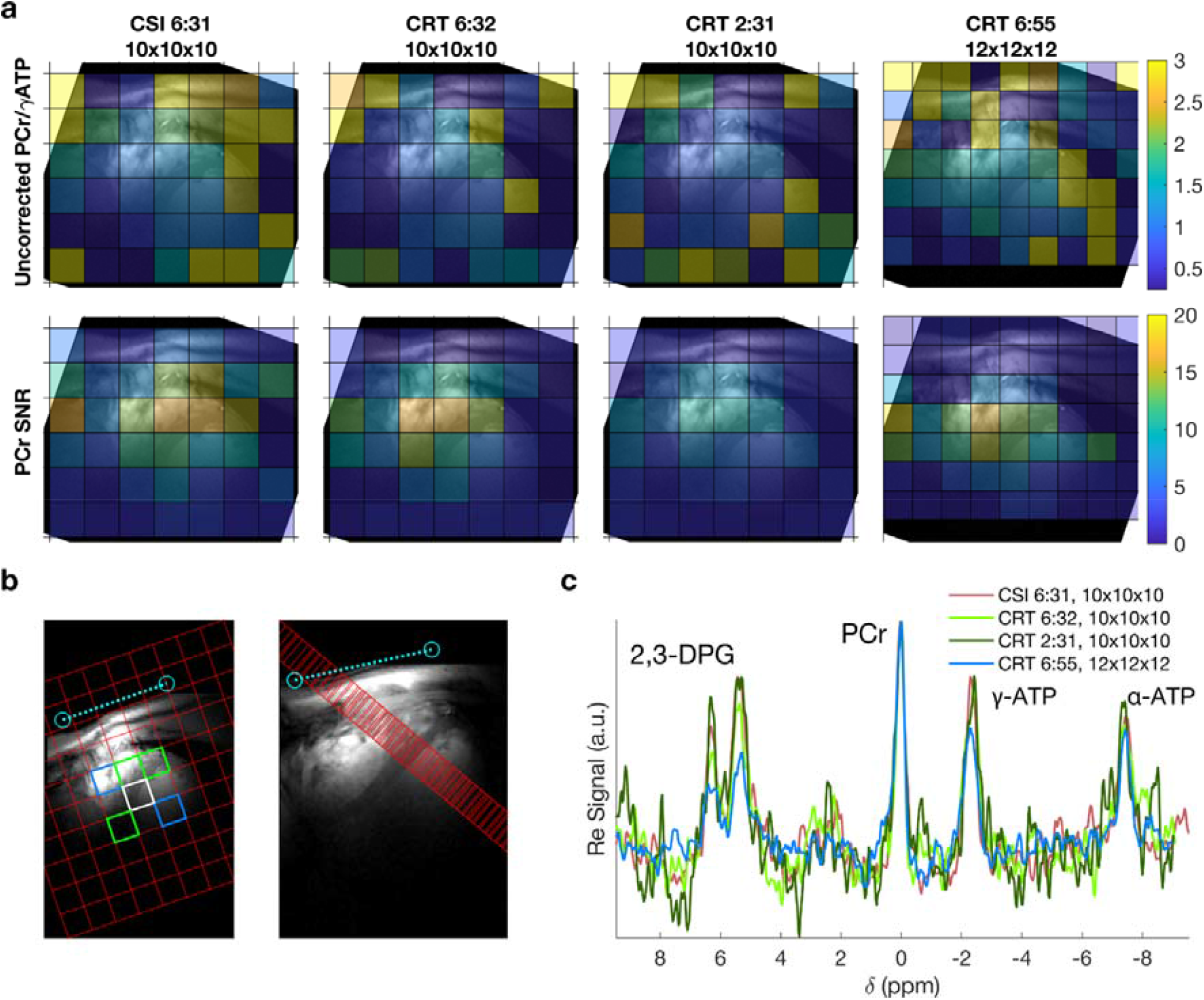
**a** Uncorrected PCr/γ-ATP ratio and PCr SNR maps of a mid-short-axis slice for four of the tested sequences overlaid on short-axis localiser images. Maps from all protocols are shown in Supporting Information Figure S1. **b** Target voxel locations overlaid on short-axis and four-chamber localiser images. Green + white = septum, blue = right and left ventricle blood pools. The intersection of one of the ^31^P coil elements with the proton image is marked in light blue. c Spectra from a single subject’s mid-septal voxel (white in **b)** for four of the tested sequences. Line colours correspond to Table 1. Spectra in c have been apodized using a 40 Hz exponential filter.

Results of the in vivo comparison are summarised in Table 1. The matched filter PCr peak SNR followed the expected relationship decreasing in line with total acquisition time and voxel volume. The 10×10×10 isotropic resolution results in a nominal voxel volume 11.5 mL and the high-resolution (12×12×12) a nominal volume of 6.7 mL. PCr SNR was 7.3 or greater for the shortest 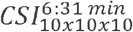 sequence, with a mean (±SD) of 9.0±6.0. A higher SNR (9.2±6.8) was measured for the high resolution 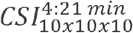 sequence. More than 73% of septal voxels could be quantified with a relative PCr/ATP CRLB < 25% for all sequences. Over 82% of voxels could be quantified for all sequences with a relative PCr/ATP CRLB < 30%.

Table 1 and Figure 6 summarise the range of PCr/ATP ratio values measured in this study. A clear dependence on slice (apical, mid, basal) is observed for uncorrected PCr/ATP values, ranging from approximately two (apical) to one (basal). Saturation and blood correction (Figure 6b) reduces this dependence, though it remains.

**Figure 6.**
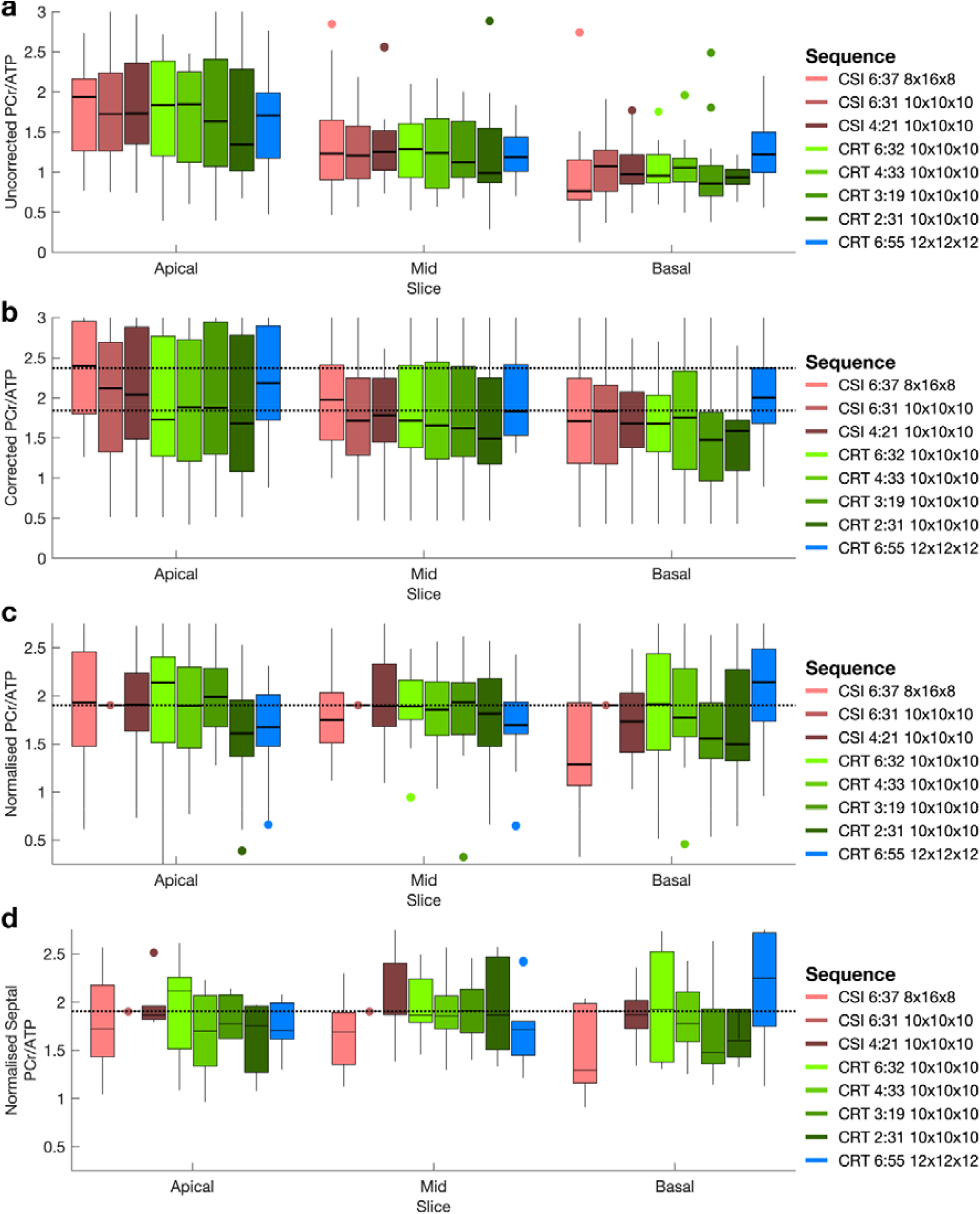
**a** Measured PCr/ATP ratios from all five subjects in target voxels (Fig 5b) from apical, mid, and basal short-axis slices in each tested sequence variant. **b** Saturation and blood corrected PCr/ATP ratios from all five subjects in target voxels (Fig 5b) from apical, mid, and basal short-axis slices in each tested sequence variant. Dashed lines show the range of measured corrected PCr/ATP ratios from Ellis et al (11). **c** PCr/ATP ratios normalised per-voxel to the values of the 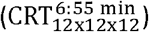 Dashed line shows normalised value scaled to mean septal value. **d** PCr/ATP ratios of just the septal voxels (green + white Fig 5b) normalised to the values of the 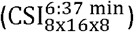 sequence. Dashed line shows normalised value.

Figure 6c&d shows values of PCr/ATP normalised to the per-voxel value measured by the 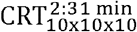 sequence, and then scaled to the mean septal PCr/ATP value. Median values measured by CRT sequences were close to the CSI values in the mid slice (Figure 6c) and mid slice septal (Figure 6d) voxels. Variance is seen to increase as acquisition time decreases. Values are less consistent in apical and basal slices. Different through plane resolutions (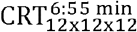 and 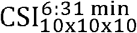) produce notably different results despite the application of blood and saturation correction.

The statistical analysis of corrected PCr/ATP values indicated that only the high resolution 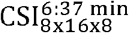 scan measured significantly different (p<0.05, Wilcoxon signed rank test) PCr/ATP distributions from the reference CSI scan.

### Denoised CRT results

All low-rank denoised spectra showed apparent denoising (Figure 7b). Across all protocols measured SNR, ignoring the effect of non-uniform variance, was 2.2 times higher than the original ‘noisy’ spectra. However, there was substantial variance between subjects in each protocol and between protocols, as shown by the large SD reported in Table 1. Denoised and corrected PCr/ATP values are reported in Table 1, and show similar mean values, though only slightly reduced standard deviations which is driven by inter-slice range. However, measured PCr/ATP ratios normalised to isotropic CSI measured PCr/ATP (Figure 7c) showed reduced variance, particularly for the intermediate duration CRT protocols. Across all denoised CRT protocols, mean denoised RMSE was 89±8% of the original noisy RMSE (compared to the values of the reference 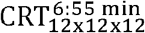 sequence).

**Figure 7.**
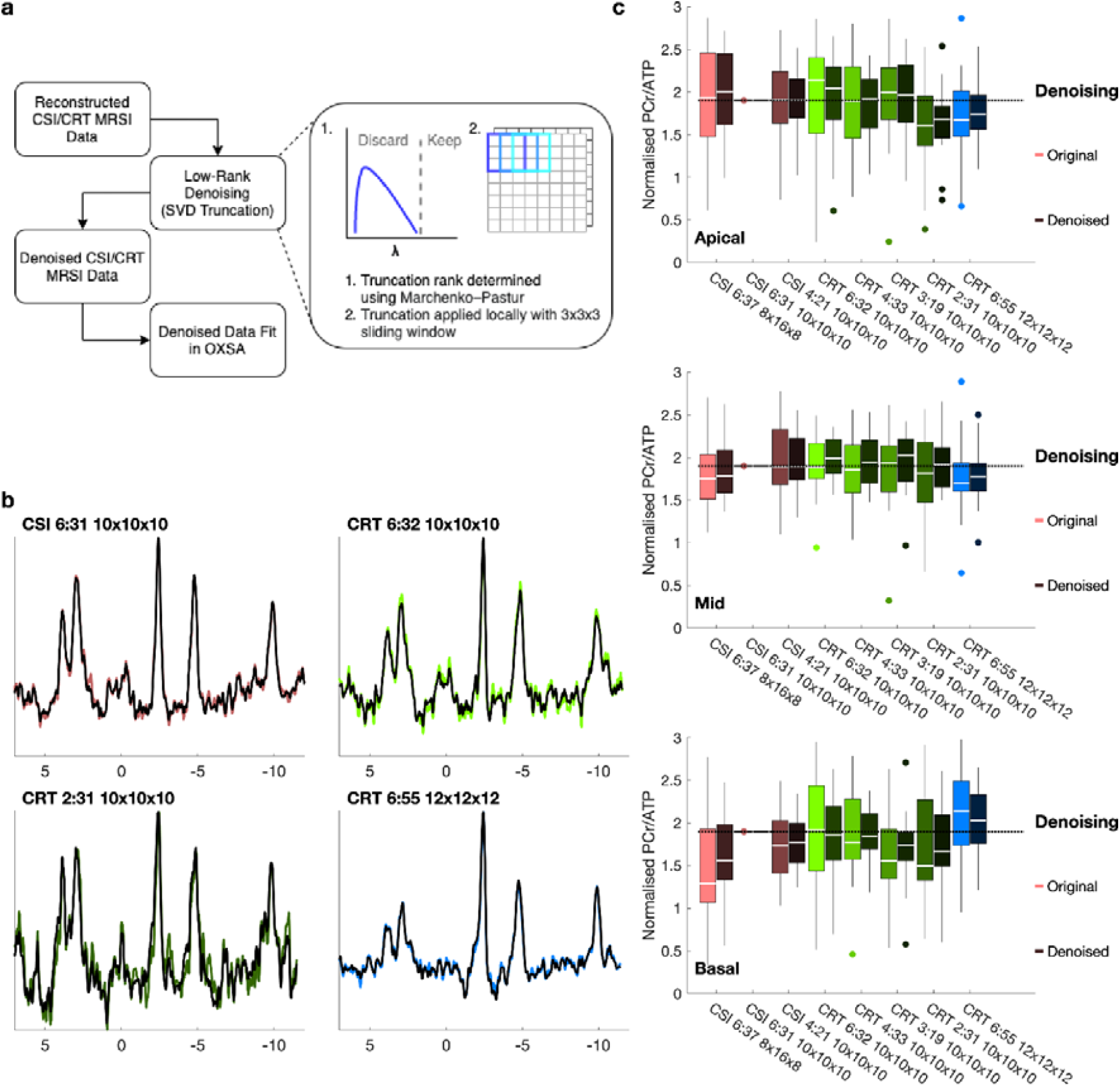
Low-rank denoising. **a** Denoising process. Reconstructed noisy CSI and CRT undergo low-rank denoising before being refit in OXSA. Automatic rank selection is used to truncate overlapping 3×3×3 patches of MRSI data, with the result being the average of the overlapping patches. **b** Denoised (dark) mid-septal spectra overlaid on original noisy data (light) from one subject and four scans (isotropic CSI, long and short CRT, and high-resolution CRT). **c** Normalised PCr/ATP ratios of original (noisy, light colour) and denoised (dark colour) voxels in apical, mid, and basal slices.

Only the 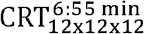 protocol was found to have a statistically different (p<0.05, Wilcoxon signed rank test) blood and saturation corrected PCr/ATP distribution from the reference, not denoised 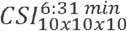.

## Discussion

Density-weighted 3D CRT MRSI has been demonstrated for fast 3D-localised cardiac ^31^P-MRS at 7T. Phantom measurements showed no loss of SNR compared to SNR optimal CSI encoding (4). In vivo PCr/ATP maps are consistent with maps generated from previously published 3D phase encoded CSI sequences. Specifically, PCr/ATP values in the inter-ventricular septum, a common region of interest, have been found to be comparable to previously published CSI sequences.

The use of a 3D density-weighted CRT sequence allows for flexibility in the acquisition time of a ^31^P-MRSI sequence, offering the ability to acquire rapid measurements without degradation of the PSF or further loss of SNR at a chosen resolution and T_R_. In this study a 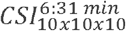 acquisition has been shown to produce good quality results not significantly different from a conventional 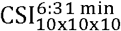 scan of identical resolution. This demonstrates a 2.63x and 1.73x reduction in scan time compared to previously described (11) and the shortest feasible Cartesian-sampled CSI sequence at matched T_R_ 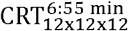. In turn, this allows for several acquisitions to be performed within a scan session, for instance during an exercise or pharmacological stress intervention (with subsequent recovery) for dynamic information about the cardiac energetics. This would be also particularly desirable for creatine kinase flux measurements, which are currently extremely long. At both 3T and 7T protocols of four scans (27) and six scans (6) respectively have been suggested, with scan times of 84 and 82 minutes. A three scan protocol has also been proposed for 3T taking approximately 70 minutes (28). Whilst the saturation transfer methods lower PCr SNR, so the same speed-ups might not be possible, use of CRT could reduce scan times from over an hour to the 30–40-minute region. One further use might be to decrease the time required for 3D resolved acquisition of cardiac Pi utilizing adiabatic excitation and long TR at 7T (29).

The implementation of the CRT sequence allows greater flexibility in the trade-off between spatial resolution and acquisition time than CSI. In this work we demonstrated a higher resolution sequence 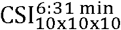, with a 6.6 mL voxel. This was acquired in 6:55 min, which is shorter than the minimum time predicted for a single average CSI sequence (8:35 min) of the same resolution. The longer CSI sequence would also suffer from SNR loss due to post-acquisition re-weighting if the same PSF was desired. In this work no density compensation was applied in the CRT sequence reconstruction, however it is likely the acquired trajectory deviates slightly from the desired density weighting function, therefore compensation might yield small improvements in PSF and SNR.

The PCr/ATP ratios measured in this study are in good agreement with those measured by Ellis et al (11), falling well within the standard deviation measured by the previous study. While the same CSI sequence, spectral processing, fitting, and saturation and blood signal contamination correction was applied here, different RF coil hardware was used. Hence, the small differences in measured PCr/ATP between the studies probably reflect the different coil transmit profiles combined with imperfect saturation correction applied. This also could explain the differences in PCr/ATP ratios measured for different slices (apical, mid, and basal) in this study. This does not interfere with the comparison of the acquisition schemes, as the same correction was applied across all variants.

Resolution was found to have a strong effect on the measured PCr/ATP ratio. Both sequences with higher 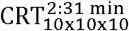 and lower 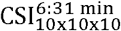 through plane resolution than the 10×10×10 isotropic resolution measured different PCr/ATP ratio distributions. The direction of the change depended on the location of the voxel being compared. This is despite standard literature saturation and blood signal contamination corrections being applied. As is evident in Figure 5c the high-resolution CRT has much lower 2,3-DPG signal in the septal voxels than the lower resolution datasets, yet blood correction does not fully account for the lower PCr/ATP measured in septal voxels of apical and mid slices. It is likely that the differing PSFs interact with the high (skeletal muscle) and low (blood or liver) PCr/ATP compartments to produce these differences.

Low-rank denoising produced a substantial denoising effect, as shown by reduced variance of PCr/ATP and lower RMSE compared to the ‘gold standard’ CSI sequence. Thus low-rank denoising has the potential to substantially mitigate the loss of SNR resulting from faster acquisitions. However, denoising will necessarily bias the measured PCr/ATP ratio and leads to non-uniform signal-dependent variance (17), so care must be taken in its use. For use in dynamic protocols with repeated sequence acquisitions, it might be possible to employ strategies to ensure similar denoising performance on each datapoint, such as rank estimation on the whole dynamic dataset.

Using CRT has some potential limitations compared to CSI. Aliasing in CRT sequences is incoherent. Therefore, in CRT datasets, aliased signal could subtly influence measured metabolite ratios in myocardial voxels without obvious visual artefacts (as is the case in CSI). This is important in the case of cardiac ^31^P-MRS where potentially contaminating tissues (skeletal muscle and liver) which contain the same metabolites at different concentrations are in close proximity. Care must also be taken with the increased susceptibility to off-isocentre distortions caused by gradient non-linearity and strong spatial blurring associated with aliased spectral peaks, if the chosen spectral bandwidth is too narrow (30). We overcame the limits on spectral bandwidth by using relatively narrow-band excitation. We are not aware of aliasing in our reconstructions.

## Conclusion

In this work we introduce a 3D density-weighted CRT sequence for rapid acquisition of ^31^P-MRSI in the human heart. The sequence is implemented on a whole-body Siemens 7T scanner. The proposed sequence can measure the PCr/ATP ratio in the human septal myocardium in 2½ min, which is 2.63x times faster than a standard CSI sequence with the same nominal voxel size of 11.5 mL. CRT can acquire high resolution data (6.7 mL voxel volume) in only 6:55 min versus the minimal 8:35 min predicted for a single-average weighted CSI, while retaining equal SNR. Low-rank denoising is particularly beneficial at these short scan times.

## Acknowledgements

WTC is supported through funding from the Wellcome Trust and the Royal Society (102584/Z/13/Z). The Wellcome Centre for Integrative Neuroimaging is supported by core funding from the Wellcome Trust (203139/Z/16/Z). BS and WB are supported by the Austrian Science Fund (FWF), [J 4124] and [P 34198] respectively. CTR and LV are funded by Sir Henry Dale Fellowships from the Wellcome Trust [098436/Z/12/B and 221805/Z/20/Z, respectively]. The support of the Slovak Grant Agencies VEGA [2/0003/20] and APVV [#19–0032] is also gratefully acknowledged. This research was supported by the NIHR Cambridge Biomedical Research Centre (BRC-1215-20014). The views expressed are those of the author(s) and not necessarily those of the NIHR or the Department of Health and Social Care. CTR and LV contributed equally to this work.

For the purpose of open access, the authors have applied a CC-BY public copyright licence to any Author Accepted Manuscript version arising from this submission.

## Supporting Information

Supporting Information.pdf – Contains Supporting Figure S1 and a detailed description of the sequence 3D density weighting implementation.

## Footnotes

1 Equation 1 also appears as Equation 4 in Reference (15) and Equation 2 in Reference (19)

## Notes

### Competing Interest Statement

The authors have declared no competing interest.

## References

1. Neubauer S. The Failing Heart — An Engine Out of Fuel. N Engl J Med 2007;356:1140–1151 doi: 10.1056/NEJMra063052.

2. Lamb HJ, Doornbos J, Hollander JA den, et al. Reproducibility of Human Cardiac 31P-NMR Spectroscopy. NMR in Biomedicine 1996;9:217–227 doi: 10.1002/(SICI)1099-1492(199608)9:5<217::AID-NBM419>3.0.CO;2-G.

3. Rodgers CT, Clarke WT, Snyder C, Vaughan JT, Neubauer S, Robson MD. Human cardiac 31P magnetic resonance spectroscopy at 7 tesla. Magnetic Resonance in Medicine 2014;72:304–315 doi: 10.1002/mrm.24922.

4. Pohmann R, von Kienlin M, Haase A. Theoretical Evaluation and Comparison of Fast Chemical Shift Imaging Methods. Journal of Magnetic Resonance 1997;129:145–160 doi: 10.1006/jmre.1997.1245.

5. Levelt E, Rodgers CT, Clarke WT, et al. Cardiac energetics, oxygenation, and perfusion during increased workload in patients with type 2 diabetes mellitus. European Heart Journal 2016;37:3461–3469 doi: 10.1093/eurheartj/ehv442.

6. Clarke WT, Robson MD, Neubauer S, Rodgers CT. Creatine kinase rate constant in the human heart measured with 3D-localization at 7 tesla. Magnetic Resonance in Medicine 2017;78:20–32 doi: 10.1002/mrm.26357.

7. Bottomley PA, Ouwerkerk R, Lee RF, Weiss RG. Four-angle saturation transfer (FAST) method for measuring creatine kinase reaction rates in vivo. Magnetic Resonance in Medicine 2002;47:850–863 doi: 10.1002/mrm.10130.

8. Weiss RG, Gerstenblith G, Bottomley PA. ATP flux through creatine kinase in the normal, stressed, and failing human heart. PNAS 2005;102:808–813 doi: 10.1073/pnas.0408962102.

9. Butterworth EJ, Evanochko WT, Pohost GM. The 31P-NMR stress test: an approach for detecting myocardial ischemia. Ann Biomed Eng 2000;28:930–933 doi: 10.1114/1.1310214.

10. Dass S, Cochlin LE, Holloway CJ, et al. Development and validation of a short 31P cardiac magnetic resonance spectroscopy protocol. Journal of Cardiovascular Magnetic Resonance 2010;12:P123 doi: 10.1186/1532-429X-12-S1-P123.

11. Ellis J, Valkovič L, Purvis LAB, Clarke WT, Rodgers CT. Reproducibility of human cardiac phosphorus MRS (31P-MRS) at 7 T. NMR in Biomedicine 2019;32:e4095 doi: 10.1002/nbm.4095.

12. Chiew M, Jiang W, Burns B, et al. Density-weighted concentric rings k-space trajectory for 1H magnetic resonance spectroscopic imaging at 7 T. NMR Biomed 2018;31 doi: 10.1002/nbm.3838.

13. Hingerl L, Bogner W, Moser P, et al. Density-weighted concentric circle trajectories for high resolution brain magnetic resonance spectroscopic imaging at 7T. Magn Reson Med 2018;79:2874–2885 doi: 10.1002/mrm.26987.

14. Hingerl L, Strasser B, Moser P, et al. Clinical High-Resolution 3D-MR Spectroscopic Imaging of the Human Brain at 7 T. Investigative Radiology 2020;55:239–248 doi: 10.1097/RLI.0000000000000626.

15. Greiser A, Kienlin M von. Efficient k-space sampling by density-weighted phase-encoding. Magnetic Resonance in Medicine 2003;50:1266–1275 doi: 10.1002/mrm.10647.

16. Nguyen HM, Peng X, Do MN, Liang Z. Denoising MR Spectroscopic Imaging Data With Low-Rank Approximations. IEEE Transactions on Biomedical Engineering 2013;60:78–89 doi: 10.1109/TBME.2012.2223466.

17. Clarke WT, Chiew M. Uncertainty in denoising of MRSI using low-rank methods. Magnetic Resonance in Medicine 2021;n/a doi: 10.1002/mrm.29018.

18. Strasser B, Chmelik M, Robinson SD, et al. Coil combination of multichannel MRSI data at 7 T: MUSICAL. NMR in Biomedicine 2013;26:1796–1805 doi: 10.1002/nbm.3019.

19. Pohmann R, Kienlin M von. Accurate phosphorus metabolite images of the human heart by 3D acquisition-weighted CSI. Magnetic Resonance in Medicine 2001;45:817–826 doi: 10.1002/mrm.1110.

20. Luo Y, Graaf RA de, DelaBarre L, Tannús A, Garwood M. BISTRO: An outer-volume suppression method that tolerates RF field inhomogeneity. Magnetic Resonance in Medicine 2001;45:1095–1102 doi: 10.1002/mrm.1144.

21. Rodgers CT, Robson MD. Coil combination for receive array spectroscopy: Are data-driven methods superior to methods using computed field maps? Magnetic Resonance in Medicine 2016;75:473–487 doi: https://doi.org/10.1002/mrm.25618.

22. Fessler JA, Sutton BP. Nonuniform fast Fourier transforms using min-max interpolation. IEEE Transactions on Signal Processing 2003;51:560–574 doi: 10.1109/TSP.2002.807005.

23. Schaller B, Paritmongkol W, Magill AW, Robson M, Rodgers CT. Quadrature 31P and single 1H dual-tune coil for cardiac 31P-MRS at 7T. In: Proc. Intl. Soc. Mag. Reson. Med. 24. Singapore; 2016. p. 4006.

24. Purvis LAB, Clarke WT, Biasiolli L, Valkovič L, Robson MD, Rodgers CT. OXSA: An open-source magnetic resonance spectroscopy analysis toolbox in MATLAB. PLoS One 2017;12:e0185356 doi: 10.1371/journal.pone.0185356.

25. Vanhamme L, van den Boogaart A, Van Huffel S. Improved Method for Accurate and Efficient Quantification of MRS Data with Use of Prior Knowledge. Journal of Magnetic Resonance 1997;129:35–43 doi: 10.1006/jmre.1997.1244.

26. Veraart J, Novikov DS, Christiaens D, Ades-aron B, Sijbers J, Fieremans E. Denoising of diffusion MRI using random matrix theory. NeuroImage 2016;142:394–406 doi: 10.1016/j.neuroimage.2016.08.016.

27. Schär M, El-Sharkawy A-MM, Weiss RG, Bottomley PA. Triple repetition time saturation transfer (TRiST) 31P spectroscopy for measuring human creatine kinase reaction kinetics. Magnetic Resonance in Medicine 2010;63:1493–1501 doi: 10.1002/mrm.22347.

28. Schär M, Gabr RE, El-Sharkawy A-MM, Steinberg A, Bottomley PA, Weiss RG. Two repetition time saturation transfer (TwiST) with spill-over correction to measure creatine kinase reaction rates in human hearts. Journal of Cardiovascular Magnetic Resonance 2015;17:70 doi: 10.1186/s12968-015-0175-4.

29. Valkovič L, Clarke WT, Schmid AI, et al. Measuring inorganic phosphate and intracellular pH in the healthy and hypertrophic cardiomyopathy hearts by in vivo 7T 31P-cardiovascular magnetic resonance spectroscopy. Journal of Cardiovascular Magnetic Resonance 2019;21:19 doi: 10.1186/s12968-019-0529-4.

30. Mayer D, Levin YS, Hurd RE, Glover GH, Spielman DM. Fast metabolic imaging of systems with sparse spectra: Application for hyperpolarized 13C imaging. Magnetic Resonance in Medicine 2006;56:932–937 doi: 10.1002/mrm.21025.

