## Supporting Information for "2½-minute 3D 7T ^31^P-MRSI of the human heart using concentric rings (CRT)"

### Sequence implementation

This supporting information details the implementation of the k-space trajectory used in the proposed 3D density weighted CRT MRSI sequence. Details of the density weighting implementation and the elliptical sampling are given.

### Density weighting

#### $K_{xy}$ weighting

Density weighting was applied in each of the three k-space dimensions. The density weighting implementation in the $k_{xy}$-plane (concentric rings) was implemented as described in Reference (Z), which describes the implementation of the original $^{1}$H sequence. The implementation was modified with a reparameterization so that the weighting function in this work (Eq [1]) matched the definition of the weighting function in References (15,19).

$$w(k) = \frac{\beta}{2} NA_{tot} \left(1 + \cos\left(2\pi \frac{k\Delta x_{\mathrm{nom}}}{\alpha}\right)\right), \qquad [1]$$

The full description of this method is given in Reference (13) in the subsection "DW-CONCEPT" of the Theory section. Further details are given in Appendix B of the same work. The methodology and code implementation was kept as close as possible to the original source code to reduce the possibility of implementation error. Here we detail the reparameterization with key equations of Reference (13) reformulated to match the weighting function used here.

Noting that $\Delta x_{\mathrm{nom}} = 1/2k_{max}$ we rewrite Eq [1] in the format of Reference (13) to express the weight as a function of $K(k)$, which translates from a regular set of k-space radii $k$, to the desired non-uniform distribution $K(k)$. Where K(k) is the needed sampling density at point $k$ in k-space.

$$w(K(k)) = \frac{\beta}{2}\left(1 + \cos\left(\pi \frac{K(k)}{\alpha k_{max}}\right)\right), \qquad [S3]$$

Equation 6 of Reference (Z) is then written

$$\frac{dK(k)}{dk} K(k) = \frac{\beta}{2}\left(1 + \cos\left(\pi \frac{K(k)}{\alpha k_{max}}\right)\right), \qquad [S4]$$

This is integrated to form an equivalent to Equation B2 in Appendix B of Reference (13) to give k(K), ther k-space position with sampling density K.

$$c_1 k(K) = \frac{\beta K^2}{4} + \beta \frac{(\alpha k_{max})^2}{2\pi^2} \cos(\pi K/\alpha k_{max}) + \beta \frac{\alpha k_{max} K}{2\pi} \sin(\pi K/\alpha k_{max}) + c_2. \qquad [S5]$$

The function *K(k)* is then approximated using an inverse power series expansion to an arbitrary order (here approximated up to 30$^{th}$ order). The inverse power series expansion was calculated using the *InverseSeries* and *Series* functions of Wolfram Mathematica (Wolfram Research Inc, Champaign, IL, USA ).

### $K_z$ weighting

The density weighting in the z-direction used a different though similar numerical approach. Both methods implemented the same weighting function, Equation [1]. Here the position of the irregularly spaced planes was calculated by first forming the numerical cumulative distribution function of Equation [1] with α = 1.61 and β=1.25, and evaluated between the limits $-\frac{\alpha}{2}, \frac{\alpha}{2}$. This was performed using the *CDF* function of Wolfram Mathematica (Wolfram Research Inc, Champaign, IL, USA ). The numerical approximation to the function $K(k_z)$, which translates from a regular set of k-space positions $k_z$, to the desired non-uniform distribution $w(k_z)$ was then approximated using a power series expansion of the inverse function to $30^{th}$ order. The power series expansion of the inverse function was calculated using the *InverseSeries* and *Series* functions of Wolfram Mathematica (Wolfram Research Inc, Champaign, IL, USA ).

The approximation $K(k_z)$ was then evaluated uniformly between 0 and 1, with the number of steps equal to the desired number of partitions. These values were then scaled by $k_{max}$ as calculated for uniform weighting. This leads to partitions non-uniformly distributed according to the desired weighting function and positioned between $-\frac{\alpha k_{max}}{2}, \frac{\alpha k_{max}}{2}$.

### Elliptical sampling

The number of concentric rings in each partition are scaled between a maximum value equal to the number of partitions and a minimum of two. The scaling is performed according to the following formula:

$$N_{rings}(i) = \left\lceil \sqrt{N_{partitions}^2 - \frac{i^2 N_{partitions}^2}{((N_{partitions}-1)/2)\text{^}2}} \right\rceil,$$

$$i = -(N_{partitions}-1)/2, \ldots, 0, \ldots, ((N_{partitions}-1)/2.$$

Where $N_{partitions}$ is the total number of partitions in the $k_z$-direction and is also equal to the maximum number of rings, and *i* is the index of the current partition. ⌈…⌉ denotes the ceiling function.

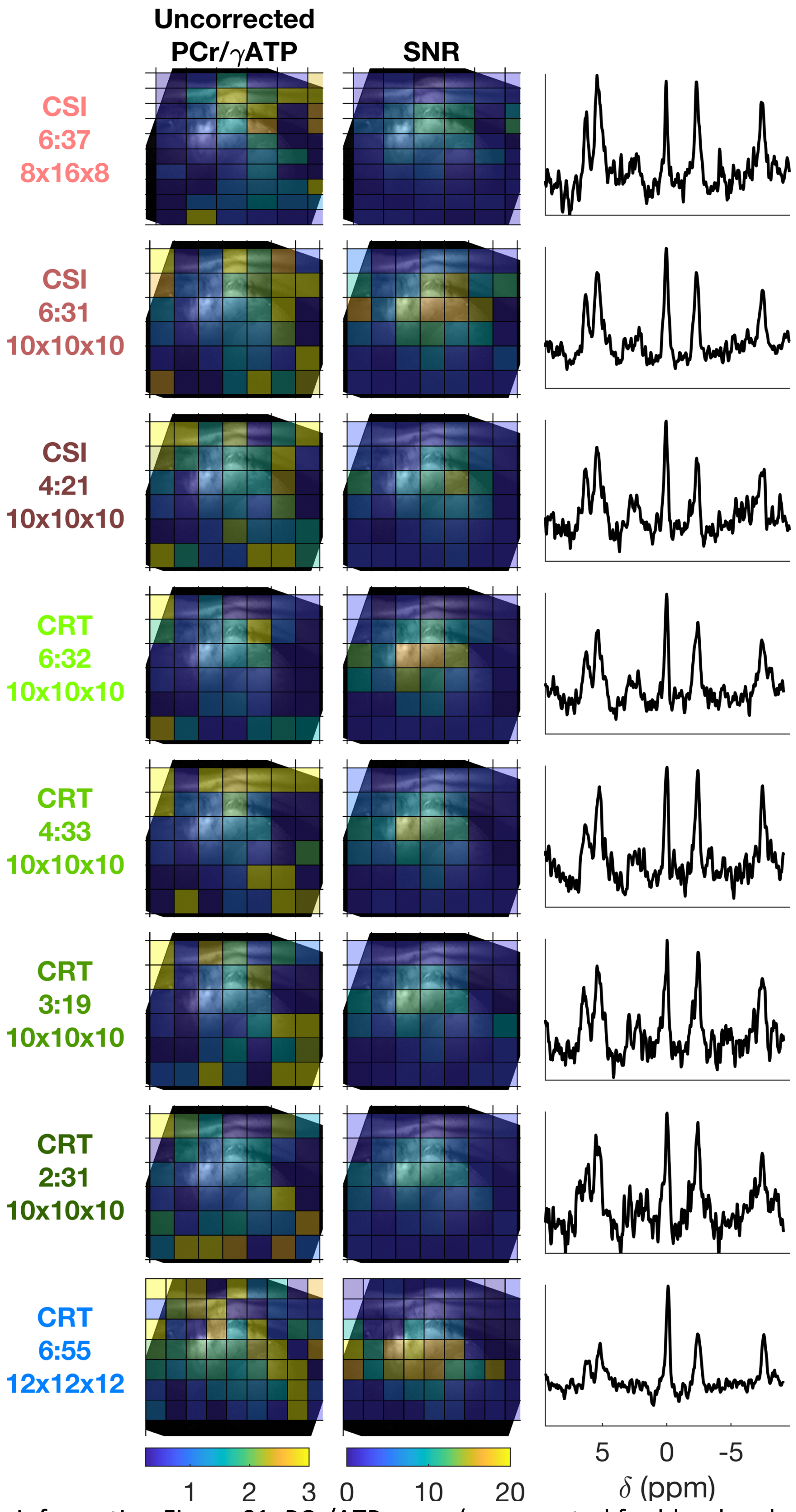


Supporting Information Figure S1: PCr/ATP maps (uncorrected for blood and saturation), SNR maps and mid-slice interventricular septal spectra for each protocol in Table 1 from a single subject.
